# scGNN+: Adapting ChatGPT for Seamless Tutorial and Code Optimization

**DOI:** 10.1101/2024.09.30.615735

**Authors:** Yi Jiang, Shuang Wang, Shaohong Feng, Cankun Wang, Weidong Wu, Xiaopei Huang, Qin Ma, Juexin Wang, Anjun Ma

## Abstract

Foundation models have transformed AI by leveraging large-scale data to efficiently perform diverse tasks, and their applications in bioinformatics are primarily focused on data-centric tasks like cell type annotation and gene expression analysis. However, their potential extends beyond data analysis, offering significant opportunities in software development and optimization, such as code refinement, tutorial generation, and advanced visualization. For example, models like OpenAI Codex can suggest optimized code snippets and generate well-documented, reproducible workflows, enhancing accessibility and reducing computational complexity. Despite these advantages, the use of foundation models for improving computational tool engineering in single-cell research remains underutilized. To address this gap, we developed scGNN+, a web-based platform that combines the power of graph neural networks with the capabilities of ChatGPT to enhance reproducibility, code optimization, and visualization. scGNN+ further simplifies the process for users by generating standardized, well-annotated code, making complex procedures more accessible to non-programmers. Additionally, ChatGPT integration allows users to create high-quality, customizable visualizations through natural language prompts, improving data interpretation and presentation. Ultimately, scGNN+ offers a user-friendly, reproducible, and optimized solution for single-cell research, leveraging the full potential of foundation models in bioinformatics. scGNN+ is publicly available at https://bmblx.bmi.osumc.edu/scgnn+.

## Introduction

Foundation models^1^ have revolutionized AI by using large-scale data to perform diverse tasks with exceptional efficiency. In bioinformatics, these models have been applied to single-cell RNA sequencing (scRNA-seq) and high-throughput sequencing data, primarily focusing on data-centric advantages such as enhanced comprehensive analysis in accuracy and scalability^2-5^. However, the potential of foundation models extends beyond basic data analysis, like cell type annotation and gene expression pattern inference; they offer significant opportunities in software development and optimization, including code refinement, tutorial generation, and advanced plotting^6,7^. For example, OpenAI’s Codex model^8^, which powers tools like GitHub Copilot, assists developers by suggesting code snippets, functions, and entire blocks of code based on user context. These capabilities enhance reproducibility by generating standardized and well-documented code that others can easily understand and replicate^9^. Moreover, foundation models can produce clear, annotated code, improving readability and making it easier for users to comprehend and modify code according to their specific needs. Algorithm optimization is another significant benefit, as these models can suggest more efficient coding practices that reduce algorithm computational demands. When it comes to plotting, foundation models can generate advanced visualization code through natural language prompts, enabling users to create high-quality, customizable charts that facilitate better data interpretation and presentation. However, the power of implementing foundation models in computational tool engineering and documentation is neglected in single-cell research.

Our in-house tool, scGNN^10^, has effectively represented gene expression and cell-cell relationships using graph neural networks. Building upon this, we developed scGNN2.0^11^, which improved gene imputation and cell clustering while reducing computational complexity. Despite these advancements, the complex procedures and high computational demands of scGNN2.0 pose challenges for non-programmers aiming to adjust the hyperparameters of models and achieve optimal results. To address these challenges, we introduce scGNN+, a web server that integrates the power of the scGNN series models with a ChatGPT^6^ assistant by leveraging the strengths of foundation models. By generating standardized and well-documented code, scGNN+ ensures that analyses can be consistently reproduced, enhancing the reliability of research findings. Beyond this, the integration of ChatGPT into scGNN+ offers several key advantages: improved readability, code optimization, and advanced plotting capabilities. The assistant provides clear, annotated code snippets, making it easier for users to understand and modify the code according to their specific needs^12^. Additionally, users can generate high-quality, customizable visualizations through natural language prompts, facilitating better data interpretation and presentation^13^.

## Prompt and standard design

To enhance user interaction and maximize the utility of scGNN+, we have designed a comprehensive tutorial prompt and code standard, as illustrated in **Fig. 1AB**. The tutorial prompt (**Fig. 1A**) guides users through every stage of the analysis by leveraging the capabilities of ChatGPT. It includes a detailed installation introduction with real-running environment information, mapping user-uploaded files to local directories, and introducing the script-running virtual environment. The prompt introduces the input data by explaining accepted file types (e.g., CSV, H5AD, Rdata) and the structure of the input data, such as the attributes stored in the AnnData object saved in H5AD files^14^. The step-by-step procedure offers an aim introduction for each pipeline step, explaining why the step is necessary and how it affects subsequent results. Alongside this, a code introduction pipeline breaks down the code used, elucidates the meaning of input parameters, and describes how these parameters influence the outcomes. The output result introduction provides detailed descriptions tailored to the specific results, including information about output directories and the data structures within saved files. Additionally, we include practice cases—both successful and failure instances—from our in-house testing. These serve as few-shot learning^15^ examples that enable ChatGPT to quickly familiarize itself with various tasks, significantly improving the accuracy of code generation and troubleshooting.

**Figure 1:**
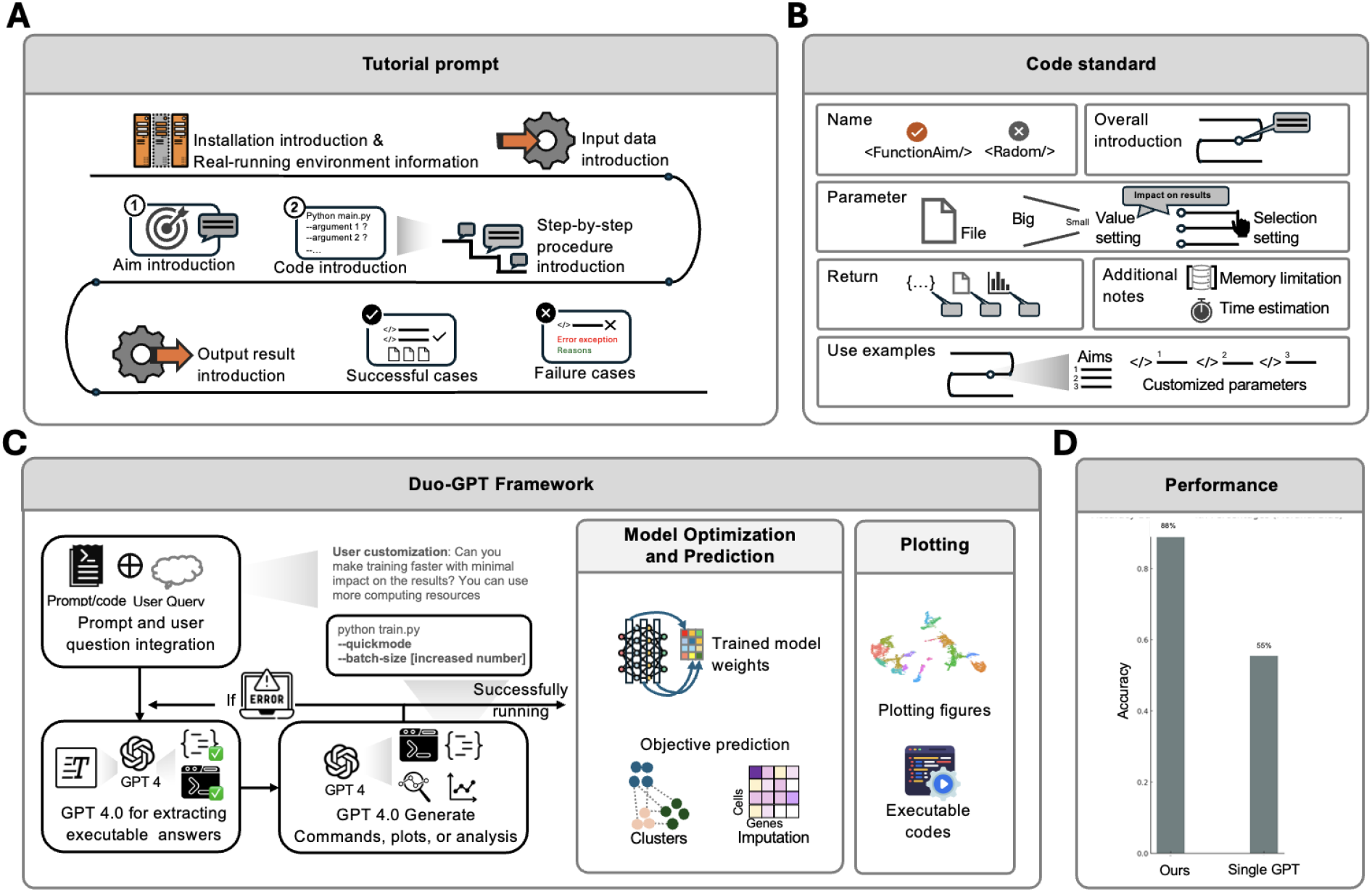
Overview of the scGNN+ framework and performance comparison. (A) Tutorial Prompt System: The tutorial prompts guide users through installation, input data formats, step-by-step procedures with code explanations, output descriptions, and practice cases. (B) Code Standards: scGNN+ designed strict code standards, including function names, descriptions, input parameters, returns, errors/exceptions, examples, and additional notes. (C) Duo-GPT framework: The scGNN+ workflow utilizes dual GPT-4 engines (Duo-GPT) to translate user queries and tutorials into executable commands and code. (D) Performance Comparison: Duo-GPT outperformed the single GPT model in code localization and customization tasks.

Furthermore, scGNN+ is developed with strict code standards (**Fig. 1B**) within the fed code for both the scGNN model and analysis procedure pipeline codes. This not only improves the accuracy of code generation but also enables GPT to provide clear explanations for the generated code. For every function used in the pipeline, we include its aim-driven name and a brief description of its purpose. We provide comprehensive details about input parameters, such as each parameter’s name, data type (e.g., string, integer, list), description, default values if applicable, and whether the parameter is optional or required. The return section specifies the data type of the output and offers an in-depth explanation of what the function returns. We also document any errors or exceptions the function might raise and under what conditions. We include sample code examples to demonstrate how the function is called and used in context, aiding users in practical implementation. Additional notes may cover constraints on memory usage limitation and time estimation. By incorporating these code standards, scGNN+ makes the code easier to understand and modify. It also enhances the accuracy of code generation and allows GPT to provide better explanations for the provided code. This further improves reproducibility and enables users to customize the functions according to their specific needs.

## Workflow design

**scGNN+** simplifies scRNA-seq analysis through an intuitive natural language interface powered by dual GPT-4 engines (Duo-GPT), shown in **Fig. 1C**. The framework integrates user queries with scGNN tutorials and code standards, translating them into large language model (LLM)^16,17^ prompts that encompass key details such as model pipelines, computational environments, hyperparameters, and output file descriptions. LLM prompt templates ensure that Duo-GPT accurately understands and generates the necessary commands and code for model training, data handling, and plotting functions. Additionally, scGNN+ allows users to define custom hyperparameters by specifying setting values and expected impacts on results within their query prompts.

Duo-GPT powers the auto-execution section, where the first GPT-4 engine generates relevant model-running commands or code, and the second GPT-4 engine validates and refines them to ensure executability within the computational environment. If errors are detected, the framework automatically corrects them and re-runs the commands or code. This seamless prompt processing enables real-time responses, reducing technical barriers for researchers working with scGNN+. Benchmarking shows that Duo-GPT outperforms single-GPT in code localization and customization tasks. Once the codes and commands are validated, scGNN+ handles the scGNN-series analysis workflow—from data preprocessing to model training—with built-in optimization steps. It incorporates key libraries like Scanpy^14^ for scRNA-seq data, Leiden clustering^18^, and UMAP^19^ for dimensionality reduction and visualization. The system iteratively trains Graph Neural Networks (GNNs)^20^, optimizing hyperparameters such as learning rates and cluster resolutions, and refining the model over multiple iterations to ensure adaptability and accuracy for tasks like cell clustering and gene imputation.

The final step in scGNN+ is visualization and interpretation. After auto-execution, the framework generates various plots—such as UMAPs, heatmaps, and cell-cell graphs—tailored to user-specified queries. Each visualization is paired with executable code, allowing users to reproduce and customize the outputs further. This integration of visualization and code enhances reproducibility and enables users to adapt the visual representations to their specific needs.

To evaluate the performance of our scGNN+ framework, we conducted a comparison between our Duo-GPT framework and a single GPT 4.0 model. We prepared several basic queries, such as “Can you help me run scGNN using the uploaded file?” and “Can you help me draw a heatmap using the scGNN imputation results?” These queries were input into scGNN+, and the process was repeated 100 times. We considered runs where the generated code was automatically executed successfully and yielded results matching the queries as correct, calculating the accuracy based on these successful executions. Similarly, we input the same questions, along with the scGNN+ tutorial and code, into a single GPT model, obtained the code, and manually executed it in the same environment as scGNN+. We then manually ran them and determined whether they ran successfully. As depicted in **Fig. 1D**, our Duo-GPT framework outperformed the single GPT model in terms of accuracy and reliability, demonstrating the advantages of our approach in code localization and customization tasks.

## Conclusion and Discussion

scGNN+ introduces a transformative approach to single-cell RNA sequencing (scRNA-seq) data analysis by addressing key challenges in the field. A significant hurdle for biologists engaging with computational tools is the reliance on programming knowledge and the complexity of pipeline instructions. Traditional web servers enable users to input data and receive output without programming but often lack the flexibility to accommodate specialized needs or adapt to novel situations. scGNN+, powered by ChatGPT, overcomes these limitations by providing a dynamic and interactive interface that not only adjusts code to meet customized requirements but also interprets results and resolves novel issues, offering a more user-centric experience.

Despite these advances, scGNN+ has certain limitations. Its extensibility is currently restricted, as it cannot automatically integrate tutorials for other tools beyond the incorporated scGNN. Additionally, the system lacks functionalities to evaluate the quality of tutorials and generate layperson-friendly guides from raw code.

To further enhance scGNN+, we aim to extend its framework to a more flexible version capable of interacting with a wide range of tools, including those for scRNA-seq and spatial transcriptomics data analysis. Another goal is to develop the ability to evaluate and score tool tutorials, providing suggestions for refinement. Finally, we aspire to enable the system to generate simplified, layperson-friendly tutorials based on input codes, broadening accessibility and ensuring that users with diverse levels of expertise can benefit from the platform.

