## Supplementary Figures for "scGNN+: Adapting ChatGPT for Seamless Tutorial and Code Optimization"

Supplementary Figure S1: scGNN+ usage cases.

Supplementary Figure S2: scGNN+ Deployment Process.

Supplementary Figure S3: Pseudo-code for tutorial prompt

Supplementary Figure S4: Pseudo-code for Duo-GPT execution process

Supplementary Information

scGNN+ usage cases

Duo-GPT Framework and Deployment of scGNN+

Supplementary Table S1: Integrated packages in scGNN+.

**Supplementary Information**

**Supplementary Figures**

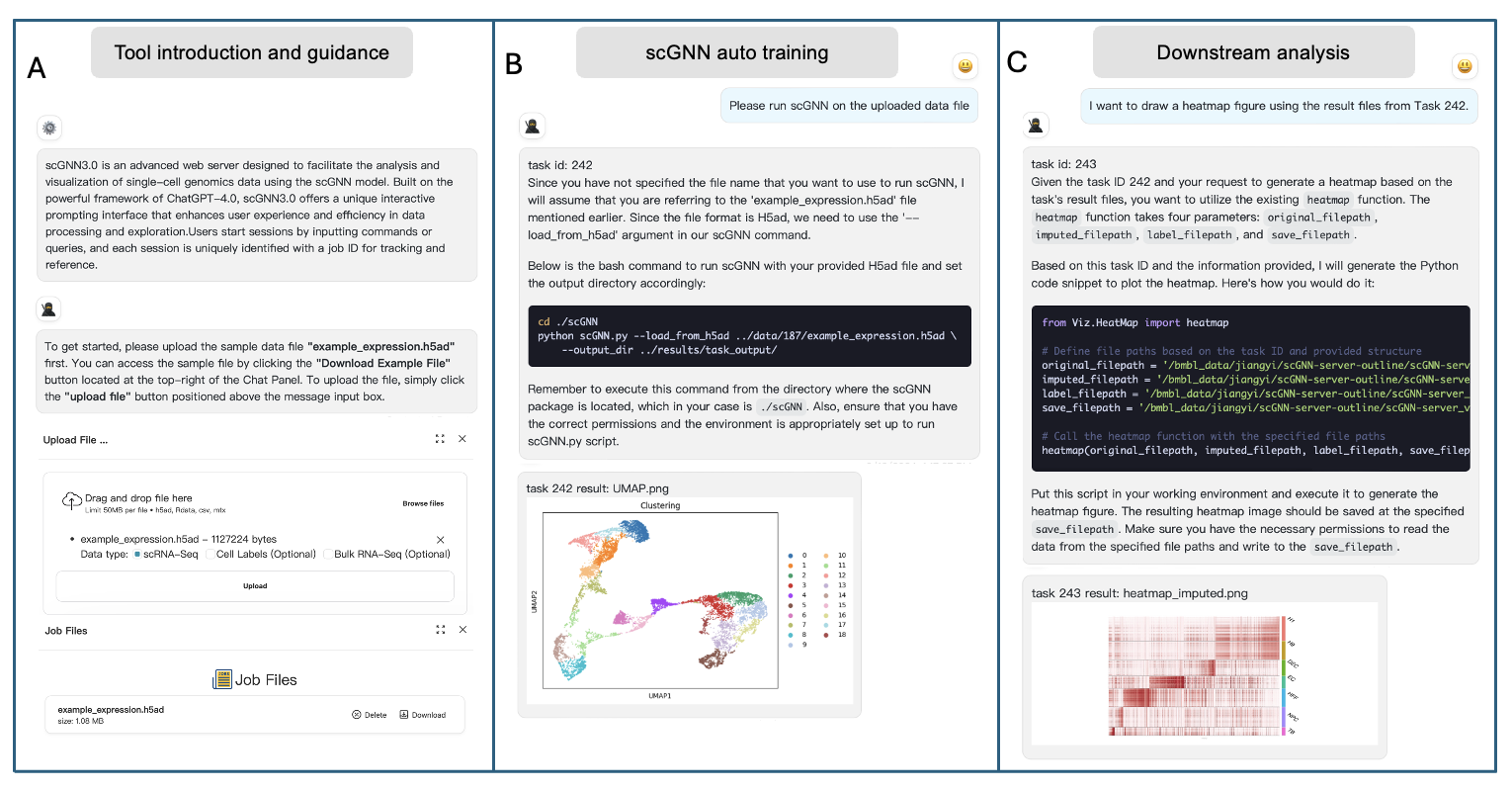

**Supplementary Figure S1: scGNN+ interactive step cases for single-cell data analysis.** This figure outlines the interactive steps in scGNN+. **(A)** shows the tool introduction and guidance where users upload their data file via a drag-and-drop interface, initiating the process. **(B)** illustrates the auto-training case where scGNN+ generates and executes a command to process the data, with results shown as a UMAP clustering plot. **(C)** demonstrates downstream analysis, where a heatmap is requested based on previous results, and scGNN+ provides Python code for heatmap generation.

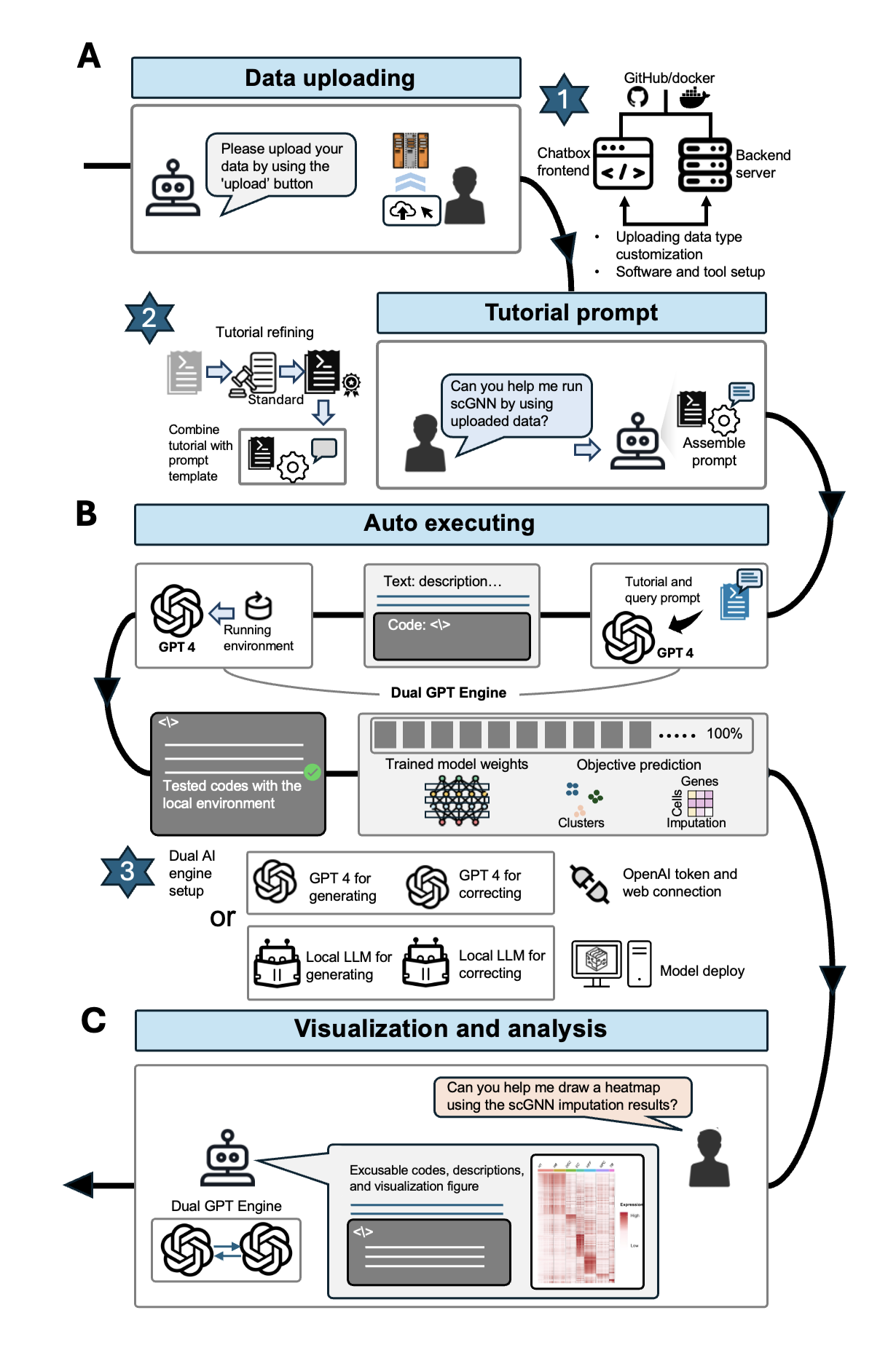

**Supplementary Figure S2: scGNN+ Deployment Process.** To deploy scGNN+, several key steps are involved to ensure a seamless user experience. The (1) step requires setting up the infrastructure necessary for scGNN’s operation. Users must clone both the front-end and back-end code from GitHub and Docker onto their local machines. After successfully cloning the repositories, users need to install and activate the required environments for both the front and back ends. Establishing communication between these two components allows users to upload their own datasets for analysis seamlessly. In the (2) step, we refined the original scGNN tutorials and code to meet the standards outlined in **Figure 1**. The tutorial and code were enhanced to integrate with the scGNN+ framework, ensuring that they provide clear guidance for users. This integrated system collects user queries and sends them to the backend, where the GPT engine processes the input and generates the necessary outputs. For the (3) step in the deployment, we implemented the Duo-GPT framework to power scGNN+'s auto-executing feature. We offer two options for setting up this framework. The first option uses OpenAI’s token to directly access the GPT-4 model, though this can be cost-prohibitive for long-term or extensive use. To address this, we provide an alternative: deploying a local large language model (LLM) on users' systems, which can mimic the same structure to form the Duo-LLM framework. This dual AI engine configuration is essential for executing tasks automatically, as it allows for both code generation and validation. Once set up, the Duo-GPT system enables scGNN+ to handle complex workflows with minimal user input, reducing technical barriers and enabling scalable bioinformatics analysis

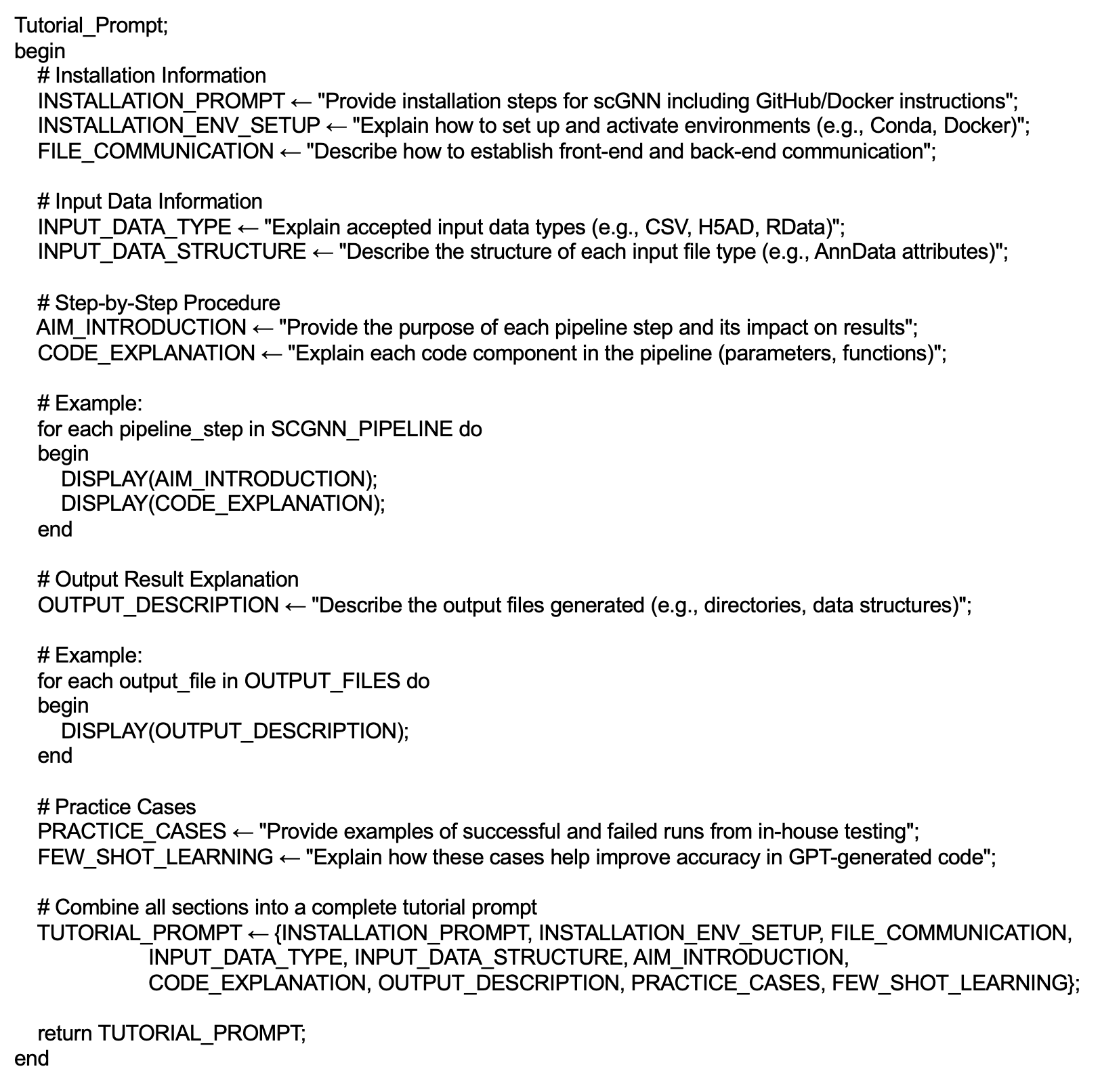

**Supplementary Figure S3: Pseudo-code for tutorial prompt** This pseudo-code adjusts the steps to represent the tutorial prompt for few-shot learning.

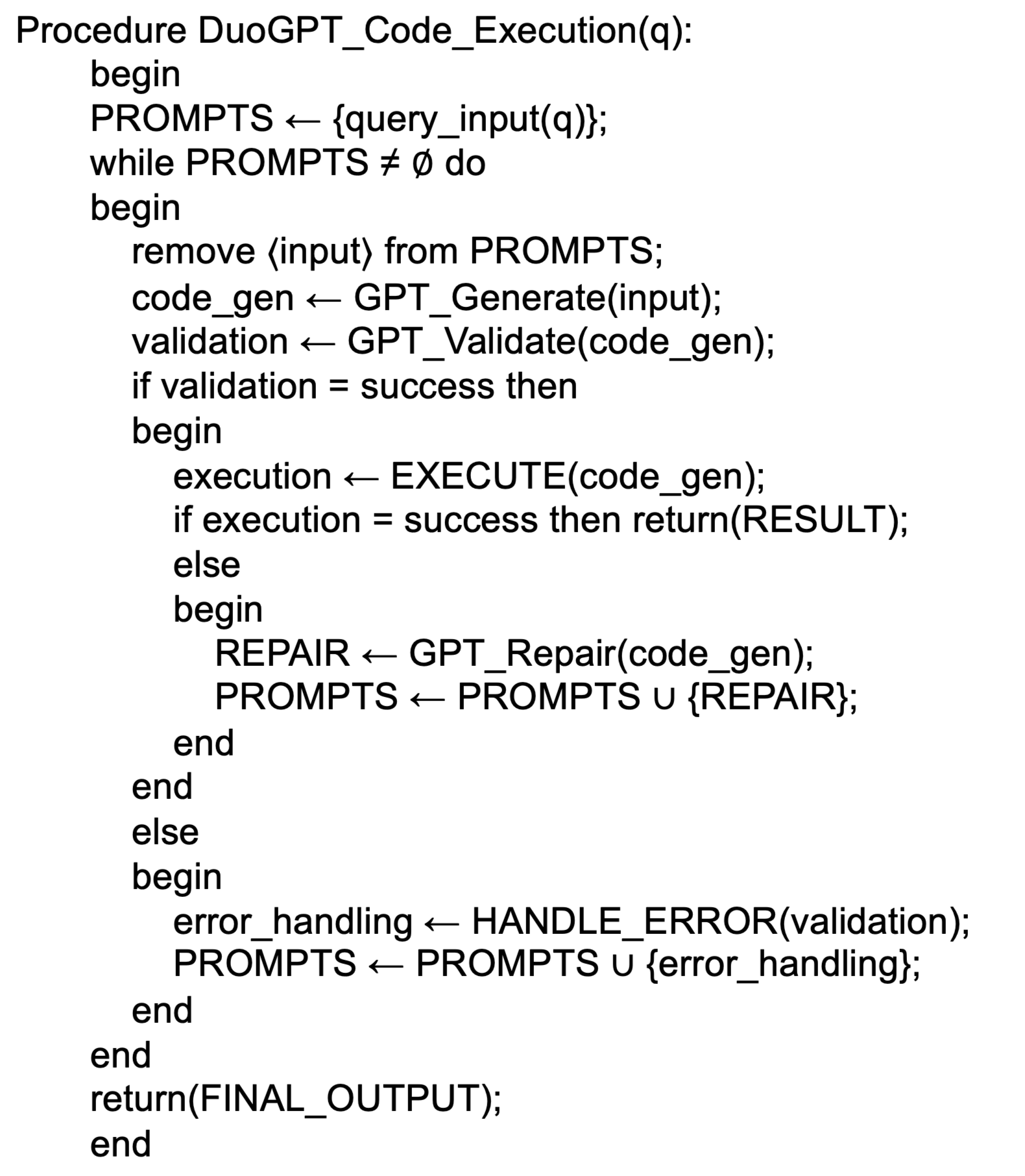

**Supplementary Figure S4: Pseudo-code for Duo-GPT execution process** This pseudo-code adjusts the steps to represent the Duo-GPT architecture for code generation and validation.

**Supplementary Information**

**scGNN+ usage cases**

Generally, scGNN+ introduces several key features that enhance both user experience and functionality. First, it integrates a chat interface, allowing users to interact intuitively by asking questions and receiving immediate feedback. Built-in usage guidance simplifies onboarding by guiding users through data processing, model training, and downstream analysis. Second, scGNN+ incorporates model auto-training, automating the process and reducing manual input to improve efficiency. Third, the platform supports comprehensive visualizations in downstream analysis, offering an end-to-end solution for single-cell data analysis.

*Tool Introduction and Guidance:* The "tool introduction and guidance" feature of scGNN+ helps users navigate the platform for single-cell transcriptomics analysis. The framework incorporates an intuitive chat interface, powered by ChatGPT 4.0, which introduces scGNN+ as an advanced platform for data processing and exploration. As part of this feature, the file management system allows users to seamlessly upload, download, and manage their data. As illustrated in Supplementary Figure S1A, users can drag and drop files or browse to select them in supported formats such as h5ad, csv, and mtx. The framework organizes these files under "Job Files," enabling users to track their uploads and manage files with options to delete or download them as needed. Each task is uniquely identified by a task ID, ensuring efficient task tracking and workflow traceability. This organized management system enhances user experience by providing easy access to files and maintaining a clear record of all completed or ongoing tasks.

*scGNN Auto Training:* Once files are uploaded, they can be used as input for model training, triggered by user queries. The "scGNN auto training" feature automates this entire process. As demonstrated in Supplementary Figure S1B, a user uploads a dataset in H5ad format and starts the scGNN model training through the chat interface. The framework assigns a task ID (task ID: 242 in this example) for easy tracking and runs the model using a predefined bash command. If the users do provide any customized preferences, the command and code will default to arguments and other pre-set options, ensuring smooth execution without manual input. After the model completes training, outputs such as clustering results are immediately generated, providing the user with visual feedback (UMAP plot). This automation minimizes user intervention, streamlining the analysis of large-scale single-cell data and optimizing the workflow for efficient data processing.

*Downstream analysis:* The "downstream analysis" feature in scGNN+ enables users to conduct various analytical tasks on the results obtained from previous stages, enhancing the interpretability of the single-cell data. In this example, shown in Supplementary Figure S1C, the user requests to generate a heatmap based on the imputed expression data from Task 242. The system assigns a new task ID (Task 243) and provides a Python code snippet to guide the user through generating the heatmap. The heatmap function requires four parameters: the original file path, imputed file path, label file path, and save file path. Once executed, the resulting heatmap is saved at the specified location, offering a clear visual representation of the data. In addition to heatmaps, scGNN+ supports a variety of other downstream analyses, such as UMAP visualization, Sankey plots, and cell-cell interaction plots. These tools allow users to explore complex relationships within their single-cell data, whether by identifying clustering patterns, visualizing gene expression distributions, or mapping interactions between cell populations. This comprehensive suite of visualization and analysis options empowers users to derive deeper insights and more meaningful interpretations from their results.

**Duo-GPT Framework and Deployment of scGNN+**

The deployment of scGNN+ follows a multi-step process, designed to ensure seamless user interaction and efficient handling of scRNA-seq analysis through natural language interfaces. Initially, Figure S2A1 shows that the infrastructure is set up by cloning both the frontend and backend code from GitHub and Docker onto the local machine. After successfully cloning the repositories, the user installs and activates the necessary environments for the front and backends. Communication is established between these components to allow the user to upload their own datasets for analysis.

For the tutorial, we began by refining the original scGNN tutorials to enhance usability. This involved providing detailed explanations for each function, clarifying result interpretations, and ensuring that all code followed standardized guidelines (Figure S2A2). Thorough code comments and structured steps improved both the clarity and integration of the tutorial with the Duo-GPT architecture. These refinements made the system more informative and user-friendly, allowing it to seamlessly interact with large language models (LLMs) like GPT-4. This combination of a refined tutorial and prompt template ensured scGNN+ could efficiently process user queries and translate them into structured, executable commands. As a result, the interaction between users and the system became more fluid, simplifying scRNA-seq analysis.

At the heart of the scGNN+ framework is the Duo-GPT engine (shown in Figure S2B), which leverages the advanced capabilities of GPT-4 to power the auto-execution feature. The system employs two GPT-4 models: Model 1 generates command-line instructions or code based on user queries, but may not always guarantee compatibility with the local environment or generate error-free code. Model 2 steps in to address these issues by refining the code, referencing error logs, successful executions, and the local environment configuration. This iterative refinement process continues until the code is successfully executed, ensuring robust error correction and optimal compatibility across different computational setups. This architecture significantly reduces the technical burden on the user and ensures that even complex code generation and validation tasks are handled with minimal manual intervention.

We provide two options for deploying the Duo-GPT framework. The first uses OpenAI’s token to directly access the GPT-4 model, though this approach may be expensive for long-term use. As an alternative, users can deploy a local large language model (LLM) using the same structure to create a Duo-LLM framework. This allows for scalable bioinformatics analysis while minimizing costs. Once either option is set up, scGNN+ automates the analysis pipeline, handling everything from code generation to validation and execution, making the entire workflow more accessible to non-programmers.

The backend of scGNN+ is powered by Flask, which manages server-side processes and task execution. Flask ensures seamless communication between the generated code and the GPT models, while managing the server load efficiently using a task queue system. This optimization supports heavy user traffic and guarantees smooth system performance even during peak usage times. On the front end, the chat-based interface, developed using Next.js, simplifies user interaction with scGNN+. Users can upload data, visualize their results, and track chat history, all through natural language commands. This intuitive interface removes the need for advanced coding skills, making it easier for users to obtain results and perform complex bioinformatics analysis.

The technical foundation (shown in Table S1) of scGNN+ is built on a combination of technologies to ensure robust functionality and scalability. Python and Rscript are used for data processing and machine learning tasks, while Flask powers the server operations. The front-end interface is built with React and Next.js, styled using HTML and CSS, to provide a responsive chat-based user experience. For artificial intelligence (AI) and machine learning, ChatGPT 4.0 facilitates natural language processing, and PyTorch supports neural network training. Anaconda (CONDA) is used for software environment management, ensuring smooth dependency handling. Nvidia hardware accelerates computation during model training and inference, and the system runs on Linux for stability, with Oracle handling backend storage and processing. This comprehensive integration of tools and frameworks allows scGNN+ to deliver an efficient, scalable, and user-friendly platform for single-cell analysis.
